# Fluorescence-based whole plant imaging and phenomics

**DOI:** 10.1101/865428

**Authors:** Stephen B. Rigoulot, Tayler M. Schimel, Jun Hyung Lee, Holly Brabazon, Kerry A. Meier, Manuel J. Schmid, Erin M. Seaberry, Magen R. Poindexter, Jessica S. Layton, Jared W. Brabazon, Jonathan A. Madajian, Michael J. Finander, John DiBenedetto, Alessandro Occhialini, Scott C. Lenaghan, C. Neal Stewart

## Abstract

Reverse genetics approaches have revolutionized plant biology and agriculture. Phenomics has the prospect of bridging plant phenotypes with genes, including transgenes, to transform agricultural fields^1^. Genetically-encoded fluorescent proteins (FPs) have transformed studies in gene expression, protein trafficking, and plant physiology. While the first instance of plant canopy imaging of green fluorescent protein (GFP) was performed over 20 years ago^2^, modern phenomics has largely ignored fluorescence as a transgene indicator despite the burgeoning FP color palette currently available to biologists^3–5^. Here we show a new platform for standoff imaging of plant canopies expressing a wide variety of FP genes in leaves. The platform, the fluorescence-inducing laser projector (FILP), uses a low-noise camera to image a scene illuminated by compact diode lasers of various colors and emission filters to phenotype transgenic plants expressing multiple constitutive or inducible FPs. Of the 20 FPs screened, we selected the top performing candidates for standoff phenomics at ≥ 3 m using FILP in a laboratory-based laser range. Included in demonstrated applications is the performance of an osmotic stress-inducible synthetic promoter selected from a high throughput library screen. While FILP has unprecedented versatility as a laboratory platform, we envisage future iterations of the system for use in automated greenhouse or even drone-fielded versions of the platform for crop screening.

## Main paper

Phenomics seeks to tightly connect genotype to phenotype across various environmental conditions^1,6^, which would enable translation of lab-based research to agricultural production and sustainability^1,7^. The various scales of phenotyping currently cover ranges from sub-micron/microscopic to satellite-based imaging of > 2000 km, with tremendous disconnect between these scales. We posit that the ‘sweet spot’ to connect genes to phenotypes as well as genomes to phenomes—for both reductionistic-mechanistic levels and ecological levels— lies at the scope of the plant canopy (meters); currently there is a technological void at this range. At the microscopic level where most basic research takes place, studies assess cellular-to-subcellular activities using state-of-the-art microscopes and molecular probes, for which innovations are numerous^3,8^. At the whole-plant-to-field level of assessment, there is tremendous potential for detecting environmental stresses on crops. The chief problem with “small scale” laboratory studies is that they are confined to tightly-controlled artificial conditions. Field experiments and radiometric models of vegetable remote sensing have the problem with long-range remote sensing is manifold: 1) ‘real-life’ systems generate data that is extremely noisy due to optical artifacts, 2) in complex environments^9^ connecting robustly-measured phenomes to genes and genomes is tenuous^1,6^, 3) incomplete illumination causes partial percent-coverage and lower leaf-area-index 4) bidirectional effects, 5) sub-pixel mixing, and 6) spatial variability on the order of a square meter in the scene landscape^10^.

In plant biology and agriculture, the most useful optical signals would be those that are completely unambiguous and occupying distinct spectral wavelengths from endogenous plant molecules. Leaf-produced compounds such as alkaloids, terpenoids, and chlorophyll produce sizable spectral ‘noise’ in plants in the form of autofluorescence^11^. In addition to avoiding spectral noise, heterologous signals should be directly tied to traits and genes. Indeed, a collection of these “ideal” spectral signatures could be stacked for multispectral signaling to expand the diversity of applications. FPs fit these criteria and can be universally imaged in plant organs. Certainly, canopy-level FP imaging is facile for UV-excitable FPs such as (near) wildtype GFP^12^ and recently-characterized GFP variants, such as those expressed in ornamental plants^13^. UV-excitable GFP can be easily imaged at the sub-meter level, e.g., seedlings and small canopies, because emission filters are not required and GFP fluorescence may be seen in the dark^5^. Previously, researchers have developed an inexpensive imaging system using blue LED arrays to excite GFP engineered into *Arabidopsis*^14^, in which dichroic filter cubes were coupled with an inexpensive camera, which could image cm-scale seedling ‘canopies.’ At the other end of the cost spectrum a portable laser-induced fluorescence imaging (LIFI) system containing a tripled Nd:YAG laser (355 nm) has been used to excite UV-excitable GFP in plants at a standoff (3 m), but this instrument was very expensive and was limited to UV-excitation because FPs excitable at 532nm were not available^10^. In order to move to higher efficiency light sources and multiple wavelengths, non-imaging techniques were explored to frequency modulate 405nm laser diodes and a fluorescence spectrometer was used to detect materials at distances greater than 2 km in field experiments with a 1 m spot size^15,16^. All current remote FP-imaging systems currently available lack flexibility with regards to imaging a variety of FPs and cannot simultaneously image multiple FPs in multiple plants at the canopy level.

Here we show the performance of a relatively inexpensive custom device (< $50,000 USD), FILP, that images plant canopies expressing various FPs at > 3 m in a laboratory setting in both constitutive and induced modalities. The flexibility of the instrument lies in the ability to select laser diodes for FP excitation and custom filters for filtering emission. Violet (400 nm), blue (465 nm) and green lasers (524 nm) were chosen in the test system to excite a range of FPs across a wide color palette (Figure 1). This laser beam is then homogenized to produce an illumination pattern that is highly uniform (flat and smooth), giving the system greater spatial resolution. Emission filters, housed in an automated filter wheel, were specifically chosen to prevent crosstalk between multiple FPs. The final major component of the system was a laboratory-grade digital camera that enabled the capture of high-resolution whole plant images. As designed, the system allows modular substitution of both diode lasers and emission filters to customize imaging of fluorescent signatures in plants, which can be accomplished at the end-user level. The goal of our study was to conduct near-simultaneous imaging of multiple, spectrally-distinct FPs at the whole canopy level in plants as a new modality of plant phenomics.

**Figure 1.**
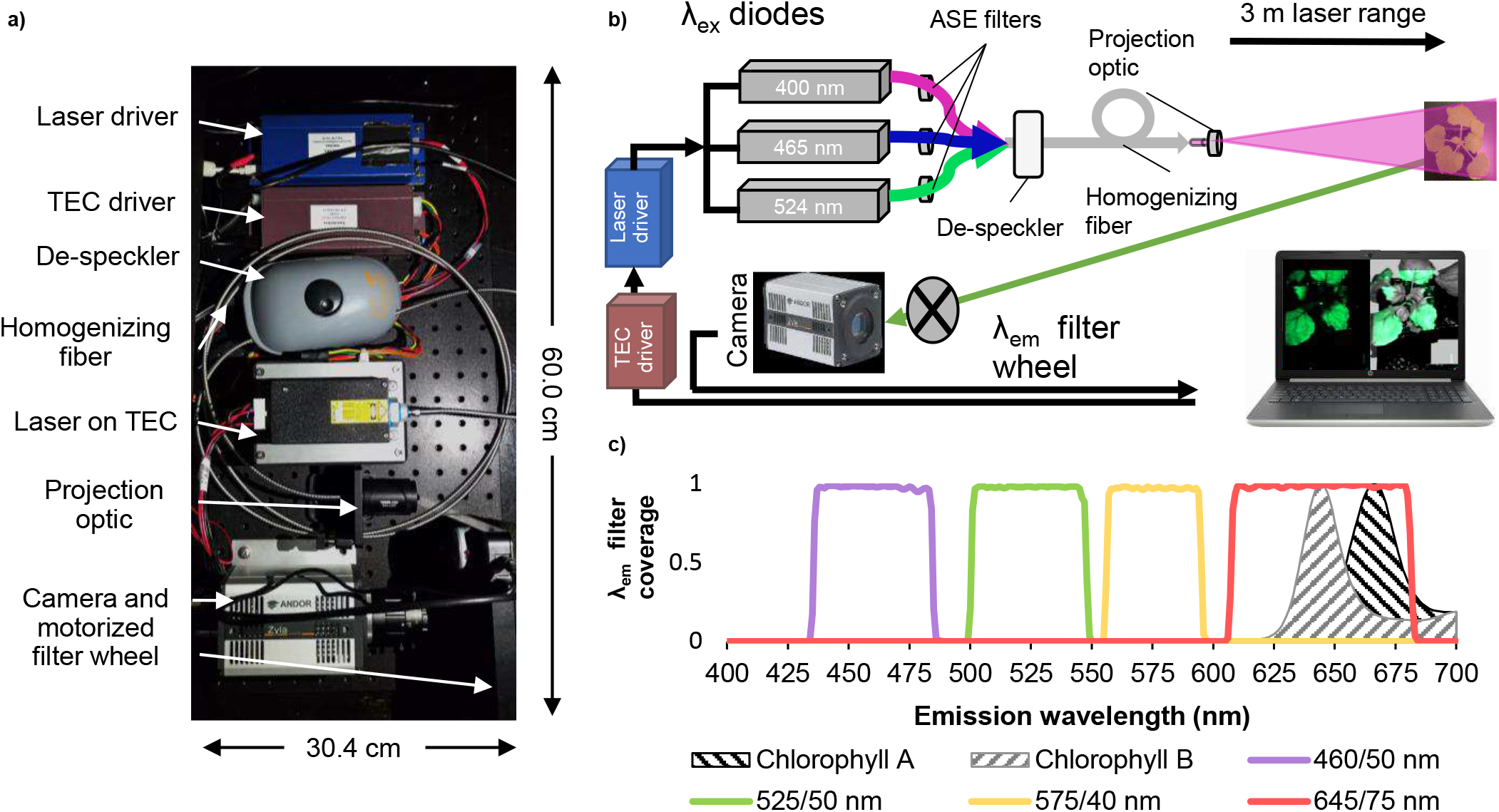
Fluoresence-inducing laser projector (FILP) characteristics. **(a)** Photograph of FILP system breadboard. **(b)** Schematic illustrating the setup of components. Abbreviations: Thermo-electric cooler (TEC); amplified spontaneous emission (ASE) **(c)** Combined line and area plot. Line plots show the wavelengths covered by the 460/50 nm, 525/50 nm, 575/40 nm and 645/75 nm notch filters, in which the first numeral is the center point and the second numeral is the breadth of the notch filter. The line plot also includes area plots which indicate the auto-fluorescence emission by chlorophylls A (excitation 614 nm) and B (excitation 435 nm) in diethyl ether.

To determine the flexibility of the FILP system, 20 FPs were characterized in agroinfiltrated *Nicotiana benthamiana*. The inherent FP excitation peaks ranged from UV (395 nm) to orange (578 nm) with emission peaks from blue (454 nm) to red (611 nm) (Table 1). The FPs also had a wide range of extinction coefficients, quantum yields and other diverse features, e.g., oligomerization states and localization tags to allow for a plethora of multi-spectral imaging schemes in plants (Table 1). Each of these FP genes were placed under the control of constitutive doubled 35S promoter in a common vector^17^ and expressed in *N. benthamiana* using either whole plant or whole leaf vacuum-agroinfiltration. Given the range of FP characteristics, many of which were suboptimal for the initial laser diodes and emission filters chosen for FILP, we were surprised that all FPs could be imaged in the plant canopies (Supplementary Figure 1). Owing to these suboptimal excitation and emission matches (FPs vs. FILP), in some cases, coupled with differences in relative brightness of FPs, there was a four-fold difference between power requirements for imaging among FPs in plants and the respective imaging channels (Supplementary Table 1). Fluorescence imaging was complemented by on-the-plant fluorescence spectroscopy measurements that modeled the three laser excitation frequencies (Supplementary Table 1). The heatmap from fluorescence spectroscopy data as well as signal-to-noise ratios of each FP emission peak, relative to a buffer infiltrated control excited at the same wavelength, was congruent with imaging results (Supplementary Table 1). Therefore, all FPs produced in plants showed similar patterns of detection via FILP consistent with quantitative fluorescence measurements.

**Table 1.** Fluorescent proteins produced in plants. Abbreviations: excitation λ (Ex), emission λ (Em); extinction coefficient (EC); quantum yield (QY); relative brightness (RB). * indicates that the EC, QY and RB data was taken from avGFP, the wild-type GFP. *Text color for mTagBFP2 and mTurquoise was changed to white to facilitate ease of reading*.

| Fluorescent protein | Donor organism | Oligomerization | Ex (nm) | Em (nm) | EC ($M^{-1}cm^{-1}$) | QY | RB | Primary reference |
| --- | --- | --- | --- | --- | --- | --- | --- | --- |
| **mTagBFP2** | *Entacmaea quadricolor* | Monomer | 399 | 454 | 50,600 | 0.6 | 32.4 | (32) |
| **mTurquoise** | *Aequorea victoria* | Monomer | 434 | 474 | 30,000 | 0.8 | 25.2 | (33) |
| **mGFP5ER\*** | *Aequorea victoria* | Monomer | 395/473 | 509 | 25,000 | 0.8 | 19.8 | (11) |
| **mEmerald** | *Aequorea victoria* | Monomer | 487 | 509 | 57,500 | 0.7 | 39.1 | (34) |
| **mT-sapphire** | *Aequorea victoria* | Monomer | 399 | 511 | 44,000 | 0.6 | 26.4 | (35) |
| **Clover** | *Aequorea victoria* | Dimer | 505 | 515 | 111,000 | 0.8 | 84.4 | (36) |
| **mVenus** | *Aequorea victoria* | Monomer | 515 | 527 | 104,000 | 0.6 | 66.6 | (37) |
| **YPet** | *Aequorea victoria* | Dimer | 517 | 530 | 104,000 | 0.8 | 80.0 | (38) |
| **PhiYFP** | *Phialidium sp.* | Dimer | 525 | 537 | 115,000 | 0.6 | 69.0 | (39) |
| **mOrange-ER** | *Discosoma sp.* | Monomer | 548 | 562 | 71,000 | 0.7 | 49.0 | (40) |
| **mKO2** | *Verrillifungia concinna* | Monomer | 551 | 565 | 63,800 | 0.6 | 39.6 | (41) |
| **LSS-mOrange** | *Discosoma sp.* | Monomer | 437 | 572 | 52,000 | 0.5 | 23.4 | (42) |
| **TurboRFP** | *Entacmaea quadricolor* | Dimer | 553 | 574 | 92,000 | 0.7 | 61.6 | (43) |
| **tdTomatoER** | *Discosoma sp.* | Tandem Dimer | 554 | 581 | 138,000 | 0.7 | 95.2 | (40) |
| **TagRFP** | *Entacmaea quadricolor* | Dimer | 555 | 584 | 100,000 | 0.5 | 48.0 | (43) |
| **mScarlet-H** | Synthetic construct | Monomer | 551 | 592 | 74,000 | 0.2 | 14.8 | (44) |
| **mRuby3** | *Entacmaea quadricolor* | Monomer | 558 | 592 | 128,000 | 0.5 | 57.6 | (45) |
| **mScarlet-I** | Synthetic construct | Monomer | 569 | 593 | 104,000 | 0.5 | 56.2 | (44) |
| **pporRFP** | *Porites porites* | Tetramer | 578 | 595 | 54,000 | 1.0 | 55.0 | (46) |
| **mBeRFP** | *Entacmaea quadricolor* | Monomer | 446 | 611 | 65,000 | 0.3 | 17.6 | (47) |

Of the 20 FPs initially screened, four of these were selected as top performers with regards to stacking and/or imaging together in canopies: mTagBFP2, mEmerald, TurboRFP and mScarlet-I (Figures 2 and 3). Multiple-FP phenomics could enable complex trait analysis. Therefore, we chose FP emissions in four distinct color bands and specific FPs that were consistently brighter than others in those color bands. Among color bands, blue-emitters seem to be the most depauperate. When designing FILP components, we purposefully matched the laser and emission filter to the optimal blue fluorescence protein spectra of mTagBFP2 (Figure 1 and Table 1). Nonetheless, both mTagBFP2 and mTurquoise could be imaged in plants using the same laser/filter combination (Figure 2 and Supplementary Figure 1). Other potential top performing FPs included both yPet and PhiYFP (Supplementary Figure 1), which would be very good choices for single FP reporters, but their emission spectra overlapped with both green and orange emitters in the 525/50 nm and 575/40 nm emission filters, which prohibited their use for simultaneous co-expression with other FPs. Top performers were also selected based on their emission peaks which were aptly spaced across the visible spectrum to facilitate robust combinatorial detection pairs. mTagBFP2 or mEmerald paired with TurboRFP or mScarlet-I were easily differentiated by FILP (Figure 2). One potential concern in imaging multiplexed FPs in the same plant cells was Förster (or fluorescence) resonance energy transfer (FRET)^18^. FRET occurs when one FP emits at the same spectrum that excites a second FP, which could potentially make dual FP detection ambiguous and confounding within the same plant. Therefore, we tested for evidence of FRET prior to selection of the four color band ‘winners’ by fluorescence spectroscopy. We observed no detectable second emission peak in the spectrophotometric measurements taken on any two co-expressed FP combinations; i.e., no FRET in our agro-infiltarted samples (Supplementary Figure 2). The initial FILP components were selected to allow for capture of optimal FP emission spectra. However, optimization of the laser light source has been facile to excite most the FPs tested using a small fraction of the total laser power available in the system. Further optimization of the emission filters may allow for the simultaneous visualization of combinations greater than two FPs per plant. In no cases did we observe any apparent laser-light damage to leaf tissues or other undesirable phenotypes when imaging plants in the FILP system.

**Figure 2.**
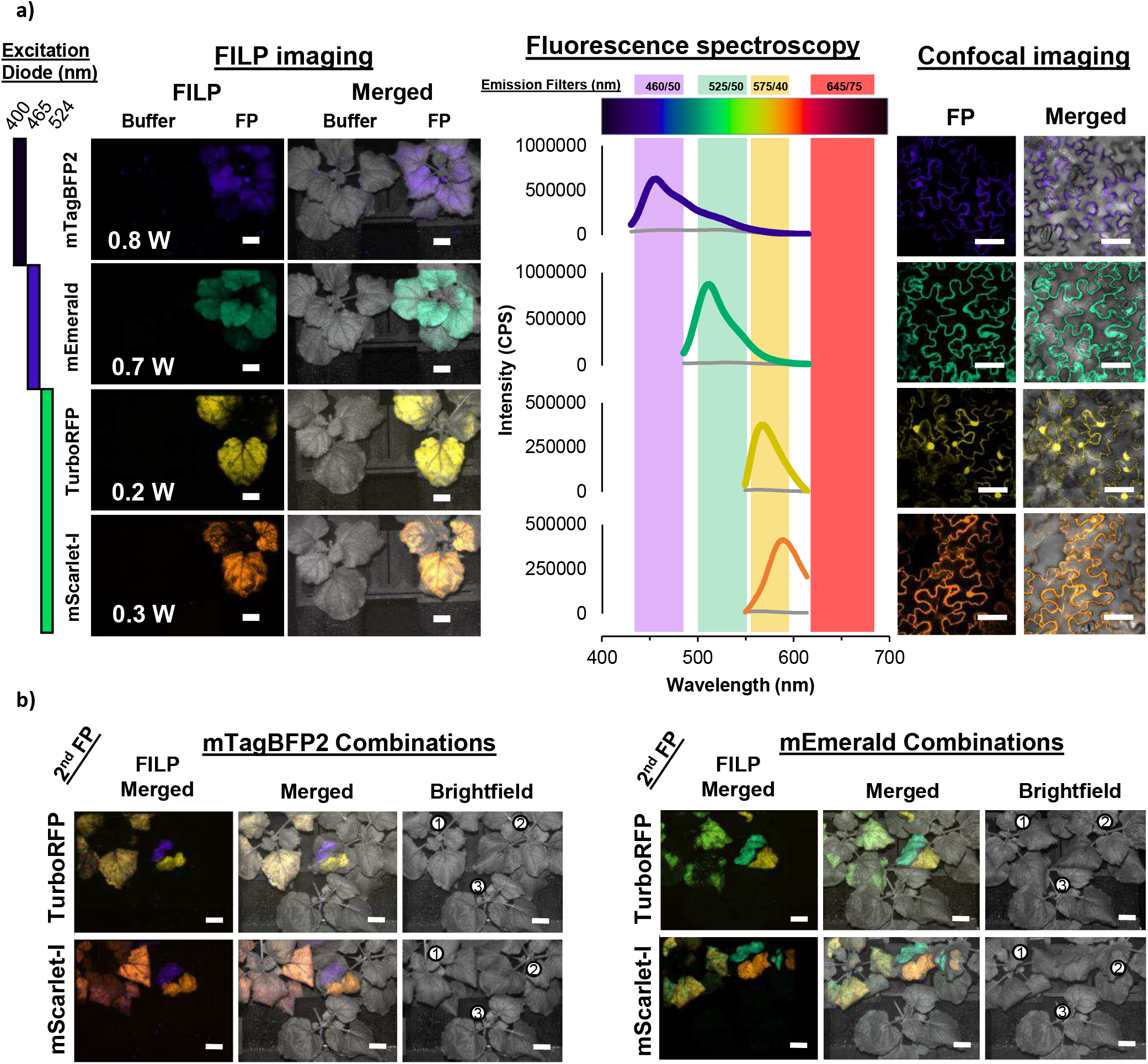
Top performing fluorescent proteins. **a)** Bars along the left side of the figure indicate which laser diode was used for excitation of each FP while the emission filter corresponds to the emission peak in the fluorolog readings. FILP images indicate laser power used in the bottom left of fluorescent images. **b)** FILP images depict four combinations using the top performing fluorescent proteins. Brightfield images for each of the combinations indicates the placement of the three plants with a circled number: 1) vacuum co-infiltrated, 2) syringe infiltrated individual FPs and 3) Buffer control. Buffer control is the same for all four combinations. Images for FILP were acquired sequentially. Laser diode and emission filter combinations are the same as those used for acquisition of single FPs. Laser power for each can be found in Supplementary Figure 2. Exposure time for FILP images was 150 ms. Scale bars for FILP images represent 2.5 cm at a detection distance of 3 m while scale bars for confocal images represent 50 µm.

**Figure 3.**
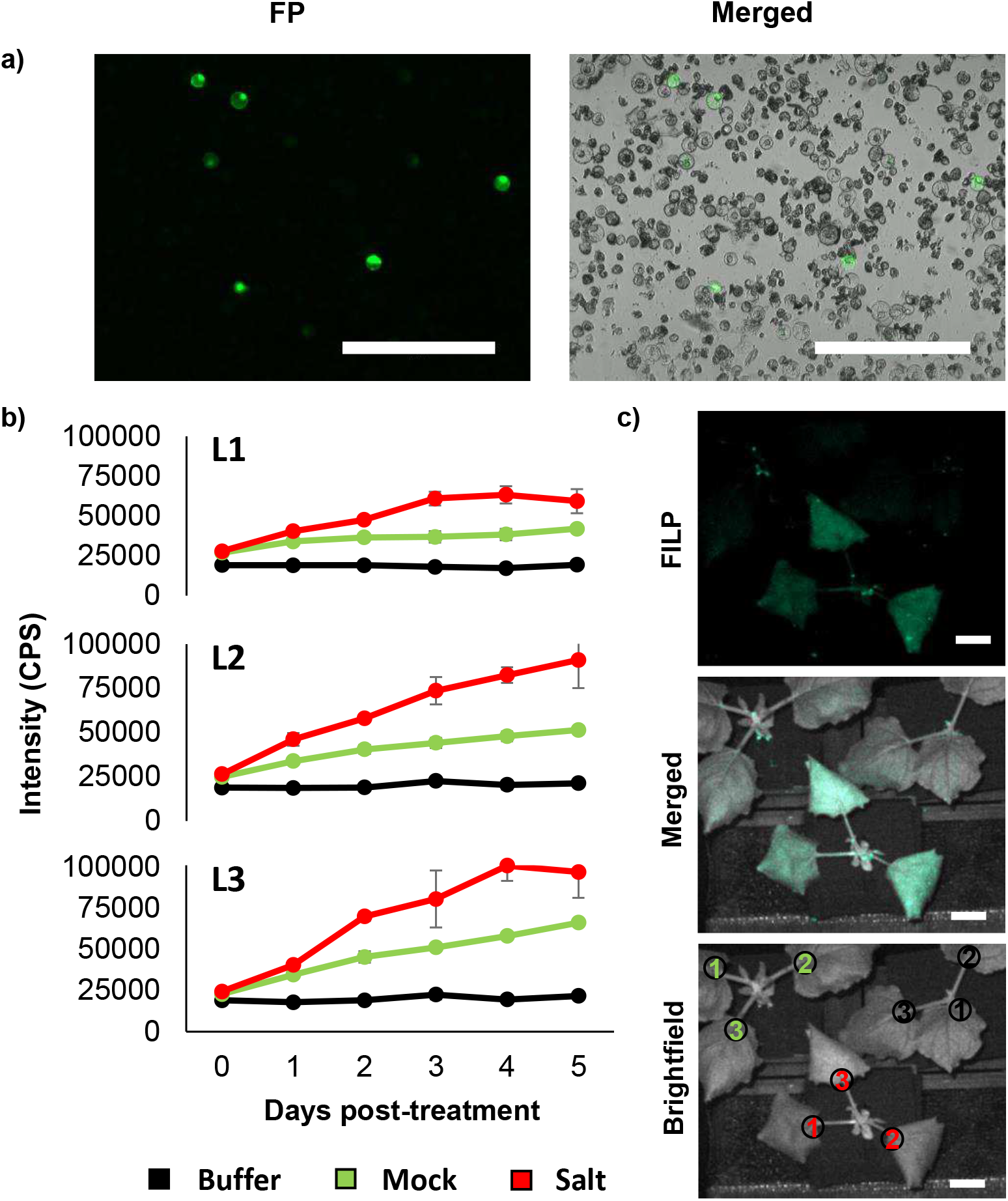
Osmotic stress-inducible promoter. **(a)** A short, synthetic promoter, ‘JL1,’ was found to be induced by osmotica in transfected protoplasts derived from a potato cell suspension culture screen based on expression of a GFP reporter. **(b)** Line plots showing fluorescence intensity measurements of the osmotic stress-inducible construct for each of the three agroinfiltrated leaves of *N. benthamiana*: leaf 1 (L1), leaf 2 (L2) and leaf 3 (L3). Error bars represent the standard deviation across four biological replicates. **(c)** FILP images of JL1 infiltrated plants 5 days post-treatment. Scale bars for FILP images represent 2.5 cm at a detection distance of 3 m. Brightfield images indicate leaf placement for each of the three treatments, buffer (black), mock (green) and salt (red), ^1^Leaf 1, ^2^Leaf 2 and ^3^Leaf 3.

The FILP system was conceived of and constructed as a phenomics device first to study plant inducibility to stresses. As an example of an ongoing study on the construction and screening of a synthetic promoter library for plants, we discovered an osmotic stress-inducible promoter in a potato protoplast screen and subsequently used FILP phenotyping to understand the promoter inducibility patterns in time and under several stimuli. Shown here is the result of one synthetic promoter (JL1) that is induced by osmotic stress (Figure 3). Five days-post osmotic stress treatment, strong inducible GFP expression could be detected in *N. benthamiana* canopies relative to control treatments. Thus, these data can be used as a first-order approximation of stand-off inducible detection in a plant model relevant to Solanaceae crops.

As plant systems biology continues to mature, arguably, phenomics may have a difficult time keeping pace with other –omics developments. One specific subset of applications of FILP-enabled phenotyping is the detection of phytosensors, which are plants engineered to detect and report environmental stimuli^19^. Phytosensors have been developed to monitor plant pathogenic bacteria to the level of field-testing^20^, but the production of clear and useful photonic signals remains challenging. In combination with advanced synthetic biology in plants^21^, especially in the area of synthetic promoters^22^ and circuits^23^, we are poised to enter the ‘golden era’ of gene-targeted phenomics. Arguably, abiotic stress detection, e.g., osmotic stress, at an early onset stage has the potential to revolutionize agricultural productivity and sustainability^24^. Our study represents the first demonstration of a ‘turn-key’ system of an osmotic stress phytosensor that can be detected optically at a stand-off. Moreover, the versatility of standoff detection using the suite of the FP color palette and the FILP phenotyping system represents an unprecedented application that clearly demonstrates the potential of FP-based phenomics in agriculture.

## Methods

### Fluorescence-inducing laser projector phenotyping system

The fluorescence-inducing laser projector (FILP) is a custom-designed instrument primarily composed of components purchased from Necsel IP, Inc. (Milpitas, CA). The Necsel components include a custom NovaLum module with three laser diodes, each emitting, respectively at 400 nm, 465 nm, and 523 nm, a Thermal Platform Developer’s Kit, an intelligent controller kit, and a homogenizing square core fiber (400 µm, 0.22 NA) used for flattening the Gaussian image produced by the lasers into a spatially uniform flat-field, albeit with residual speckle from the laser coherence. This speckle was reduced by passing the fiber through a commercially-available aquarium pump to vibrate the fiber faster than the camera’s exposure time, thereby smoothing and de-speckling the resulting image. Individual amplified spontaneous emission (ASE) filters were hand cut and mounted onto each laser diode to limit the transmitted wavelength. As such, a 405 ±20 nm filter (ZET405/20X, Chroma Technology Corp., Bellow Falls, VT) was mounted onto the 400 nm diode, a 460 ±36 nm filter (ET465/36 nm, Chroma Technology Corp.) was mounted onto the 465 nm diode, and a 524 ± 24 nm (FF01-524/24, IDEX Health & Science, LLC., Rochester, NY) was mounted onto the 523 nm diode. After assembly, the maximum power output of the complete Necsel system was 1.14 W @ 400 nm, 1.36 W @ 465 nm, and 1.45 W @ 523 nm. Further, control of the current (from the intelligent controller) allowed linear control over the laser power with R^2^ values > 0.99 (Supplementary Figure 3). To shape the beam for imaging whole plants, a projector lens (63-714, Edmund Optics, Barrington, NJ) was placed 6 mm in front of the homogenizing fiber to form a 20 cm^2^ imaging square at a distance of 3 m.

Images of plants at ≥ 3 m was achieved using an Andor Zyla 5.5 sCMOS camera equipped with a 50 mm focal length lens (86-574, Edmund Optics). The camera was mounted to the same breadboard as the laser system, ensuring that the excitation and emission distances were identical. Control of the camera resolution, exposure time, and image acquisition was achieved using the free open-source software µManager. A five-position motorized filter wheel with USB control (84-889, Edmund Optics) was mounted between the camera lens and the sample to enable collection of images for specific wavelengths pertaining to the target fluorescent protein synthesized by plants. In the current version of FILP, three 50 mm emission filters were loaded onto the emission wheel: 460 ± 50 nm (ET460/50 nm), 525 ± 50 nm (ET525/50 nm), and 575 ± 40 nm (ET575/40 nm) (Chroma Technology Corp.).

To ensure user safety of the system and provide complete darkness for sampling, a custom laser range, 0.61 m × 0.91 m × 3.7 m, was assembled around the entire system using 25 mm construction rails and black hardboard (XE25 & TB4, ThorLabs, Newton, NJ). Plants were placed inside the enclosure through a door fabricated at the back of the enclosure using the same materials. Finally, a magnetic interlock was used to ensure that the class IV lasers could be operated as a class I system where the lasers were immediately and automatically turned-off if the door was opened.

### Plant expression vectors for constitutive expression of fluorescent protein genes

The DNA coding sequences for the 20 fluorescent protein (FP) genes listed in Table 1 were mobilized into Invitrogen pENTR /D-TOPO cloning vectors (Invitrogen, Carlsbad, CA). Following colony PCR and validation by sequencing, the FP coding sequences were each recombined into pMDC32 35S expression vectors^17^ via the LR Clonase reaction (Invitrogen, Carlsbad, CA). FPs were subsequently re-sequenced prior to transformation into *Agrobacterium tumefaciens* strain LBA4404.

### Synthetic promoter screening

To identify candidate osmotic-stress inducible plant promoters, a library of synthetic promoters, totaling > 2000 constructs, was screened in potato protoplasts (Stewart et al., unpublished data). Transformed protoplasts were observed for GFP expression using an EVOS M7000 imaging system (ThermoFisher Scientific, Waltham, MA) equipped with a GFP filter (excitation: 470/22 nm, emission: 510/42 nm), and protoplasts were scored as positive or negative for induction. Promoters identified in the protoplast screen were then characterized in leaves by agroinfiltration assays in *N. benthamiana*.

For the osmotic stress treatment, each pot was watered with 100 ml of NaCl solution (250 mM) 48 hr after agroinfiltration, followed by withholding water for 5 d to partial wilt stage. The mock treatment consisted of 100 ml tap water applied every two days to each plant. Three biological replicates were used, and the experiments were repeated three times. Fluorescence spectroscopy measurements were taken immediately prior to NaCl treatment and then repeated every day for 5 d. FILP images were taken on the final day of the experiment. Leaves not previously measured by the fluorescence spectrometer were removed, including old leaves and new growth that had arisen since vacuum infiltration. The mEmerald reporter was excited using the 400 nm laser diode and observed using the 525/50 nm filter. Laser wattage was 0.8 W and the exposure time was 150 ms.

### Vacuum agroinfiltration of *N. benthamiana*

*Agrobacterium* was infiltrated according to Rigoulot et. al. (2019)^25^ with modification. *Agrobacterium tumefaciens* strain LBA4404 was used for the infiltration of all fluorescent protein constructs. Colony PCR was used to determine transformation of *Agrobacterium*. *Agrobacterium* was then grown from colonies overnight in 10 ml YEP media with rifampicin (50 mg/L) and kanamycin (50 mg/L) selection at 28°C shaking. This culture was used as the seed culture for a 125 ml culture also grown overnight under the same conditions. *Agrobacterium* was resuspended in injection media (10 mM MES, 10 mM MgCl_2_ and 100 µM acetosyringone) to an OD600 value of 0.8 and this *Agrobacterium* solution was incubated for 3 hr at room temperature prior to infiltration. Four-week-old *N. benthamiana* plants grown under long day conditions at 23°C were used for infiltration experiments. Vacuum infiltration of *N. benthamiana* plants were performed using a modified Nalgene vacuum desiccator (Cole Parmer, Vernon Hills, IL). The sidearm on the base was blocked using the PTFE cap (provided with purchase) and secured in place with parafilm. The stopcock on the desiccator lid was removed and PVC tubing was retro-fitted with a 1-5 ml pipette tip to connect the benchtop vacuum port with the vacuum desiccator. *N. benthamiana* plants were inverted and all aboveground tissues were submerged into a Magenta™ Ga-7 Plant Culture Box (Fisher Scientific, Catalog No. 50-255-176) filled with the *Agrobacterium* suspension. The Magenta box was housed inside of a Styrofoam support ring that was cut to the size of a large glass container to prevent the movement of the vessel during application and release of the vacuum. With the modified pipette tip end of the hose securely inserted into the vacuum desiccator lid, the vacuum was applied in 1 min intervals. This was repeated 3 times to achieve thorough infiltration of leaf tissue indicated by visible saturation of the leaf with the *Agrobacterium* solution. The vacuum provided by the benchtop vacuum port was measured to be −84 kpa or −12 psi. After infiltration, the plants were rinsed in a beaker of DI water and then allowed to dry at room temperature (Supplementary Video 1). The plants were then returned to the growth room until fluorolog readings, FILP and confocal imaging were taken 72 hr post infiltration.

For co-infiltrated plants, *Agrobacterium* solution was adjusted for both FP gene constructs to an OD600 of 1.6, then FP constructs were mixed 1:1 and vacuum infiltration was conducted as previously described. Syringe infiltration of *N. benthamiana* leaves was conducted as described in Rigoulot *et al.* (2019)^25^ with *Agrobacterium* solution at an OD600 of 0.8.

### Fluorescence spectroscopy of leaves

Prior to FILP measurements, the targeted spectral characteristics of the youngest, fully-expanded leaf of each plant was quantified using scanning fluorescence spectroscopy (Fluorolog®-3, obin Yvon and Glen Spectra, Edison, NJ, USA) using emission spectral acquisition by the FluorEssence Software (HORIBA Scientific, version 3.8.0.60). On-the-leaf fluorescence was measured using a fiber optic probe as described previously^26^. Excitation wavelengths matched Necsel laser diode wavelengths at 400 nm, 465 nm, and 523 nm with a slit width of 5 nm. Emission wavelengths were scanned from 415-615 nm, 480-615 nm, and 540-615 nm for the respective excitation wavelengths in increments of 1 nm. Leaves of the same developmental age were infiltrated with buffer as a negative control and measured for background fluorescence.

Fluorescence spectroscopy data was handled using custom software: the Fluorologger Shiny app, coded in R^27,28^. The graphic user interface allows for the visualization and normalization of fluorolog data and the app is currently available on github at github.com/jaredbrabazon/Fluorologger. Output files from the Fluorolog (.dat format) were input into the application and with user input the data were normalized according to methods described in Millwood et al. (2003)^26^. A detailed user guide is available at the link provided.

The heatmap in Supplementary Table 1 was created by recording the fold change difference between an individual fluorescent proteins peak emission as described on FPBase.org and the background emission at this same point taken by the fluorolog (signal to noise).

### Confocal microscopy

The same leaf tissue analyzed by fluorescence spectroscopy (Fluorolog) was imaged using an Olympus FV1200 confocal microscope (Olympus, Center Valley, PA, USA). Diodes lasers (405, 440, 473, 559 and 635-nm lasers) along with conventional Argon and HeNe (R) lasers were used to image the investigated fluorescent proteins with excitation (Ex) and emission (Em) spectra indicated in Table 1. Single confocal images are shown in the manuscript. The manufacturer’s Olympus FV10-ASW Viewer software Ver.4.2a (Olympus, Center Valley, PA, USA) and the ImageJ^27^ analysis software (version 1.41o) were used to acquire and process confocal images, respectively.

### Image processing

Assembly of FILP and confocal microscope images was done using the ImageJ^27^ analysis software (version 1.41o). Color determination for each fluorescent protein was done using the Wolfram Demonstration Project (https://demonstrations.wolfram.com/), Colors of the Visible Spectrum plugin. Using the Adobe color space option, peak emission wavelengths were used to query for RGB values. These values, representing a percentage, were multiplied by the maximum value for the R,G or B decimal code (255). The resulting values were then used to establish look up tables (LUT) for the ImageJ^29^ software. Images are input into the ImageJ software independently. Adjustments to brightness and contrast were applied uniformly across images if necessary. Using the images to stack function, FILP or confocal fluorescent and bright field (BF) images were overlaid. After using the composite image function and selecting the color option of the channels tool, pseudo coloring is applied to a selected image. Presets include the gray which can be used for the brightfield image as well as blue, green, red, etc. For more specific color palettes, a unique LUT was generated. By default, ImageJ applies different colors to the different channels (images) and these were changed using the channels tool color option. Images were exported as .tiff files for the construction of figures. We provide an example of the Wolfram player determination of R, G, B values, the conversion of these values to generate individual LUTs, the table of RGB decimal values for all FPs and a visual walkthrough of pseudo coloring (Supplementary Figure 4).

### Preparation of potato cell suspension

The preparation of potato cell suspension used for synthetic promoter screening of protoplasts was adapted from a previously described method by Sajid and Aftab (2016)^30^. Solanum tuberosum cv. ‘Desireé’ was propagated by nodal explants into propagation media [4.33g/L MS salts (Phytotech M524, PhytoTech Labs, Lenexa, KS), 25 g/L sucrose, 100 mg/L myo-inositol, 0.17 g/L sodium phosphate monobasic, 0.44 g/L calcium chloride dihydrate, 0.4 mg/L thiamine HCL, 5 ml/L “complete vitamin stock” (for 100 ml; 40 mg glycine, 10 mg nicotinic acid, 10 mg pyridoxine HCL, 10 mg thiamine HCL), 3 g/L phytagel, pH 5.8, 1 ml/L MS (Phytotech M557) vitamins) in Magenta GA7 vessels under 16-h day, 8-h night fluorescent light conditions at room temperature (23°C). Sterile leaf explants, cut into 1-2 cm squares, were taken from propagates and callus was induced on callus induction (CI) media [4.33 g/L MS salts, 20 g/L sucrose, 2 g/L gelzan (solid media), pH 5.8, 1 ml/L MS vitamins, 4 mg/L 2,4-D (2,4-dichlorophenoxyacetic acid)]. Callus was transferred to fresh CI plates every 2-3 weeks. After 4-5 weeks on CI media, approximately 2 g of green, friable callus was used to inoculate 20 ml of liquid CI media and grown on a platform shaker at 120-140 rpm for 7 d. The suspension was then filtered through a 425 µm sieve and transferred to a new 125 ml flask. After the filtered cells settled, 15 ml of liquid CI media was removed, replenished with fresh CI media and the suspension was grown again on a platform shaker at 120-140 rpm. After 7 d, an additional 30 ml of fresh liquid CI media was added to the flask and allowed to grow for another week. The cell suspension was maintained every 5-7 d by sub-culturing approximately 15 ml of the filtered suspension culture into 30 ml of fresh media. Cells were periodically filtered through a 425µm sieve to maintain consistency.

### Protoplast transfection

Potato protoplasts were isolated from cell suspension culture 3 d post-subculture. Five milliliters of packed cell volume was digested by a cell wall-digesting enzyme solution [0.4 M mannitol, 20 mM MES (pH 5.7), 20 mM KCl, 10 mM CaCl2, 1% (w/v) bovine serum albumin (BSA), 5 mM β-mercaptoethanol, 4.4% (v/v) Rohamet CL, 4% (v/v) Rohapect, 0.6% (v/v) Rohapect UF] in the dark at room temperature for 2 h with gentle shaking. After two washes with wash buffer [0.45 M mannitol, 10 mM CaCl2] protoplasts were filtered through a 40 µm nylon mesh cell strainer (Fisher Scientific, Hampton, NH) and intact cells were purified using a 23% sucrose gradient centrifugation. Protoplasts were then suspended in a MMg solution [0.4 M mannitol, 15 mM MgCl2, 4 mM MES] to a concentration of 2 × 105 protoplasts/ml. For protoplast transfection assay, PEG-mediated transfection was conducted according to Yoo et al. (2007)^31^ with modification. Ten microliters of plasmid (1 µg/µl) was added to 100 µl of protoplast suspension. After adding 110 µl of 40% PEG solution [40% (w/v) PEG-4000 (Sigma), 0.8 M mannitol, 1 M CaCl2], the mixture was incubated for 15 min at room temperature. The reaction was stopped by adding 440 µl of W5 solution [2 mM MES, 154 mM NaCl, 125 mM CaCl2, 5 mM KCl] and the protoplasts were centrifuged at 100 × g for 1 min, and then suspended in 200 µl of WI solution [0.5 M mannitol, 4 mM MES, 20 mM KCl] for incubation in the dark for 48 h.

Transformed protoplasts were observed for GFP expression using an EVOS M7000 imaging system (ThermoFisher Scientific) equipped with a GFP filter (excitation 470/22 nm, emission 510/42 nm).

### Plant stand

A custom plant stand (“FILP’s castle”) to secure potted plants being imaged in the laser range was fabricated using 3D printing. The stand tilts potted plant samples 75 degrees forward to allow for maximum foliar exposure while using the laser system. The stand accommodates three square (7.6 cm) pots that are grown in an 18 cell flat (59-3080, Griffin Greenhouse Supplies, Inc., Tewksbury, MA). Designs for 3D printing are available upon request. The face piece for plant stand was printed on a Lulzbot Taz 3D printer using nGen Copolyester with 30% infill. To create the PVC support stand, PVC pipe with approximately 22.23 mm outer diameter was cut into three 15.24 cm pieces and two 7.62 cm pieces. The weighted end piece was created by fitting each of the 7.62 cm pieces of PVC tubing into an PVC elbow and joining them in the center using a tee piece. After orienting the openings in the same direction, they were secured in place by silicone adhesive. This piece was then filled with sand and plugged by additional silicone. The weighted end piece was then attached to the openings on the back of the 3D printed face using the remaining 15.24 cm PVC pieces.

## Data Availability

Original or processed .tif image files for all FILP and confocal experiments are available from the corresponding authors upon request.

---

**Supplementary Figure 1.**
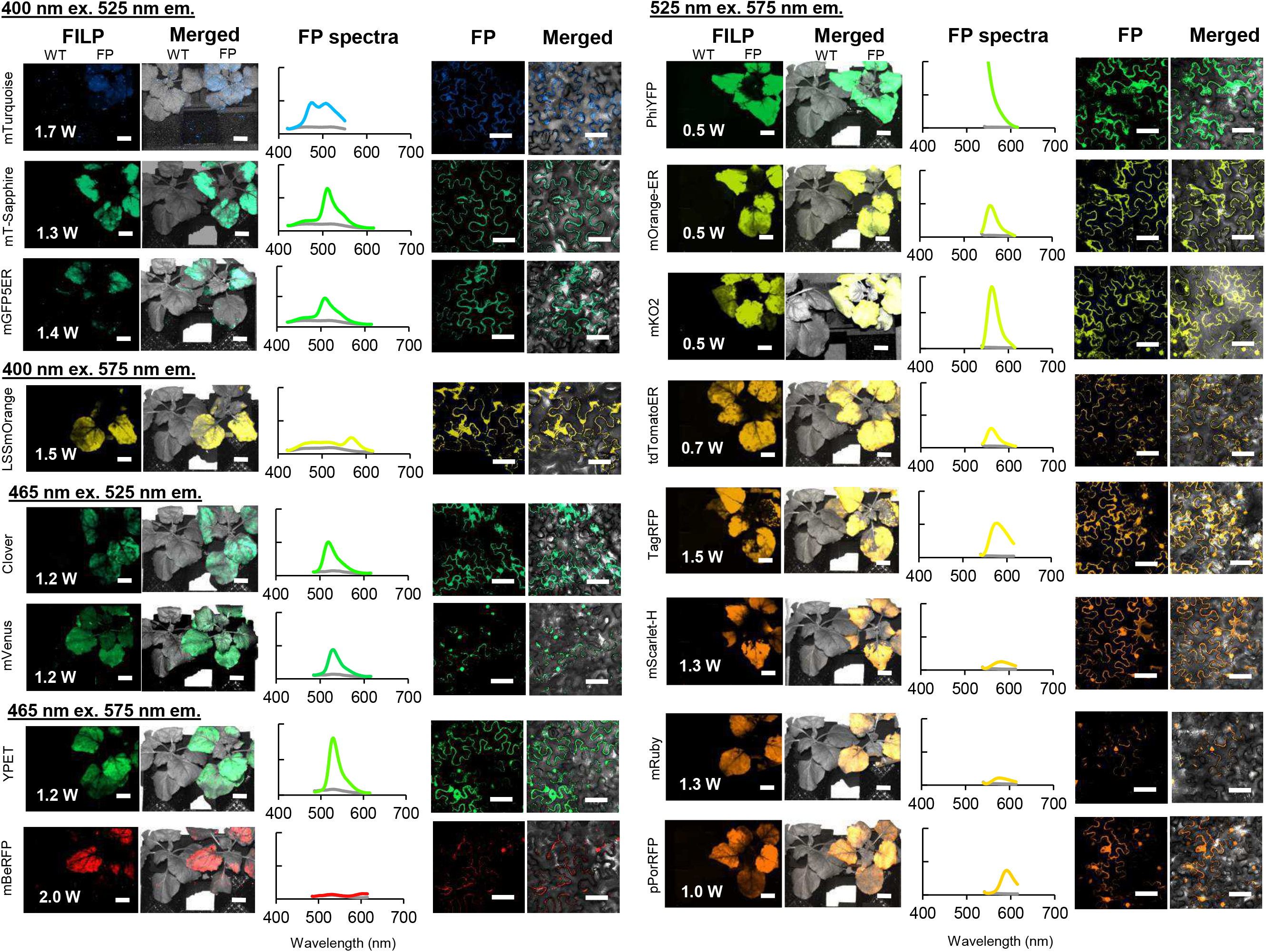
Fluorescent proteins successfully observed by the FILP system. Fluorescence spectroscopy measurements and confocal imaging included for verification. Exposure time for FILP imaging can be seen in Supplementary Table 1. Y-axis for all plots is scaled to 5 × 10^5^ CPS except for PhiYFP which is scaled to 1.0 × 10^6^ CPS. Scale bars for FILP images represent 2.5 cm at a detection distance of 3 m while scale bars for confocal images represent 50 μm.

**Supplementary Table 1.** Heatmap of laser power required for the detection of fluorescent proteins. Laser diode power and emission filter combinations are scored from no signal (none) to the highest signal to noise ratio. Heatmap of fluorescence spectroscopy measurements show fluorescence when compared to buffer infiltrated control at each fluorescent protein peak emission at 400 nm, 465 nm, and 524 nm excitation wavelengths. Diagonal line fill indicates that the excitation wavelengths exceed the reported peak emission value and therefore the fluorescence intensity for the peak emission cannot be calculated. Data acquired for this figure and Supplementary Figure 1 used an earlier version of µManager, which did not support the full capabilities of the Andor camera. As such, a higher laser voltage was required to collect this data, when compared to the data collected in the main text with a newer version of µManager.

|  |  |  |  | FILP imaging and measurements |  |  |  |  |  | Signal to noise measurements from fluorescence spectroscopy |  |  |  |
| --- | --- | --- | --- | --- | --- | --- | --- | --- | --- | --- | --- | --- | --- |
|  | Peak maxima |  | Exposure (ms) | Ex. 400 nm |  |  | Ex. 465 nm |  | Ex. 524 nm |  | Ex. 400 nm | Ex. 465 nm | Ex. 524 nm |
|  | Ex (nm) | Em (nm) |  | 460/50 | 525/50 | 575/50 | 525/50 | 575/50 | 460/50 | 575/50 |  |  |  |
| mTagBFP2 | 399 | 454 | 500 | 1.4 W | 1.4 W |  |  |  |  |  |  |  |  |
| mTurquoise | 434 | 474 | 500 | 1.7 W | 1.7 W | 1.7 W | 1.4 W | 1.4 W |  |  |  |  |  |
| mGFP5er | 395 | 509 | 500 | 1.4 W | 1.4 W |  |  |  |  |  |  |  |  |
| mEmerald | 487 | 509 | 500 | 1.4 W | 1.4 W | 1.4 W | 1.4 W | 1.4 W |  |  |  |  |  |
| mT-sapphire | 399 | 511 | 500 |  | 1.3 W | 1.3 W |  |  |  |  |  |  |  |
| Clover | 505 | 515 | 500 |  | 1.5 W | 1.5 W | 1.5 W |  |  |  |  |  |  |
| mVenus | 515 | 527 | 500 |  | 1.5 W | 1.5 W | 1.2 W | 1.2 W |  |  |  |  |  |
| yPET | 517 | 530 | 500 |  |  |  | 1.2 W | 1.2 W |  | 0.5 W |  |  |  |
| PhiYFP | 525 | 537 | 500 |  |  |  | 1.4 W | 1.4 W |  | 0.5 W |  |  |  |
| mOrange-ER | 548 | 562 | 500 |  | 1.4 W | 1.4 W |  | 2.0 W |  | 0.5 W |  |  |  |
| mKO2 | 551 | 565 | 500 |  |  |  |  |  |  | 0.5 W |  |  |  |
| LSS-mOrange | 437 | 572 | 500 |  | 1.5 W | 1.5 W |  | 1.5 W |  |  |  |  |  |
| TurboRFP | 553 | 574 | 500 |  | 1.3 W | 1.3 W |  | 2.0 W |  | 0.5 W |  |  |  |
| tdTomatoER | 554 | 581 | 500 |  | 1.8 W |  |  | 2.0 W |  | 0.7 W |  |  |  |
| TagRFP | 555 | 584 | 200 |  |  |  |  | 2.0 W |  | 1.0 W |  |  |  |
| mScarlet-H | 551 | 592 | 200 |  |  |  |  |  |  | 1.3 W |  |  |  |
| mRuby3 | 558 | 592 | 200 |  |  |  |  |  |  | 1.5 W |  |  |  |
| mScarlet-I | 569 | 593 | 200 |  |  |  |  | 1.5 W |  | 0.5 W |  |  |  |
| pporRFP | 578 | 595 | 200 |  |  |  |  | 1.5 W |  | 1.0 W |  |  |  |
| mBeRFP | 446 | 611 | 200 |  |  |  |  | 2.0 W |  |  |  |  |  |

Signal to noise ratio
None
Low
Medium
High
No Change
Log2 Fold-change
Highest

**Supplementary Figure 2.**
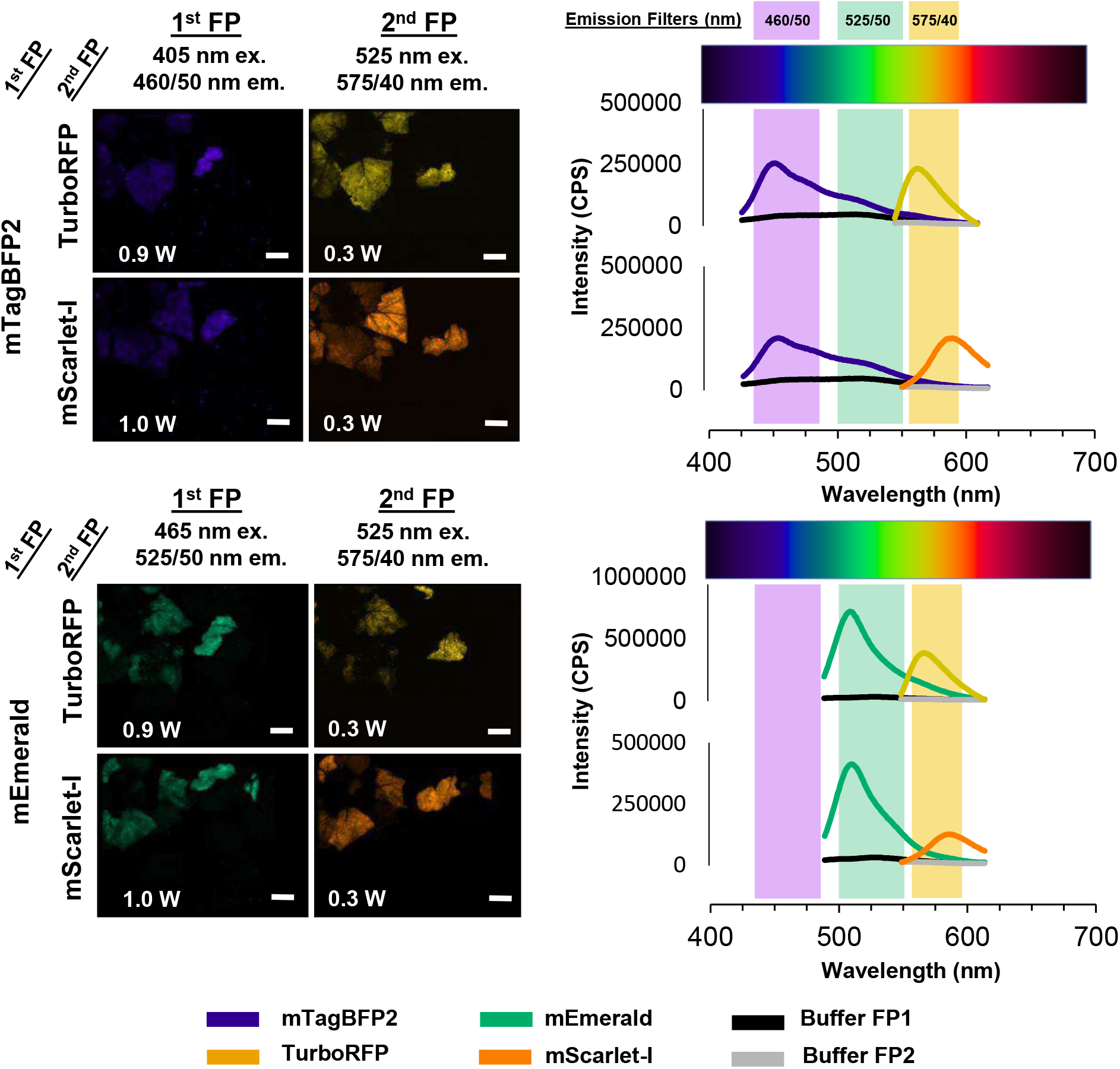
Fluorescence spectroscopy measurements of co-expressed fluorescent proteins. Individual channels of co-expressed FPs for combinations described in Figure 2. Exposure time for FILP images was 150 ms. Line plots correspond to fluorescence emission measurements taken on vacuum co-infiltrated plants. Black lines designate the buffer control plant reading for the 1^st^ FP and the buffer for the 2^nd^ FP is in grey. Scale bars for FILP images represent 2.5 cm at a detection distance of 3 m.

**Supplementary Figure 3.**
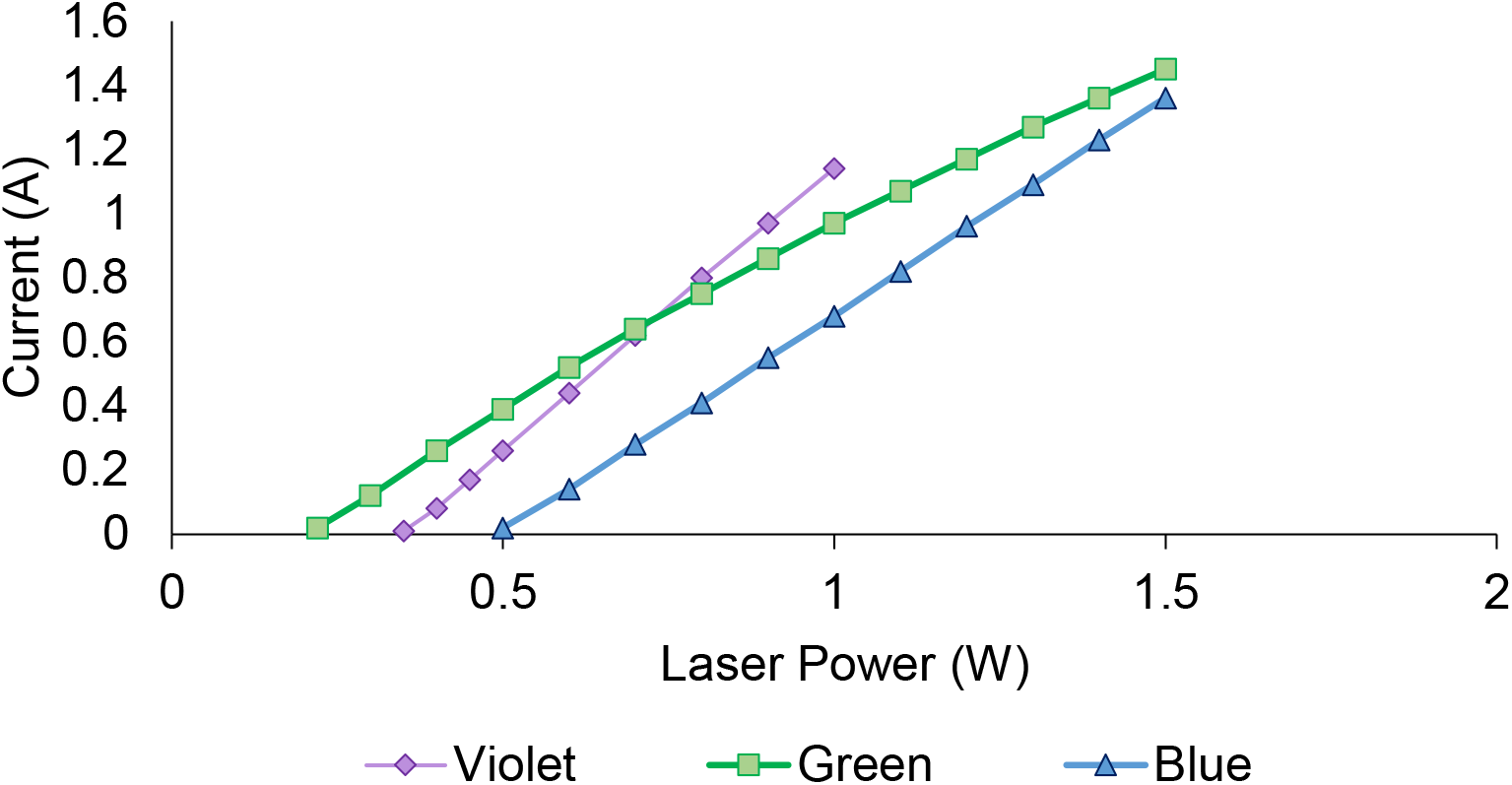
Current vs. laser power. Controlled by the Necsel Intelligent Controller.

**Supplementary Figure 4.**
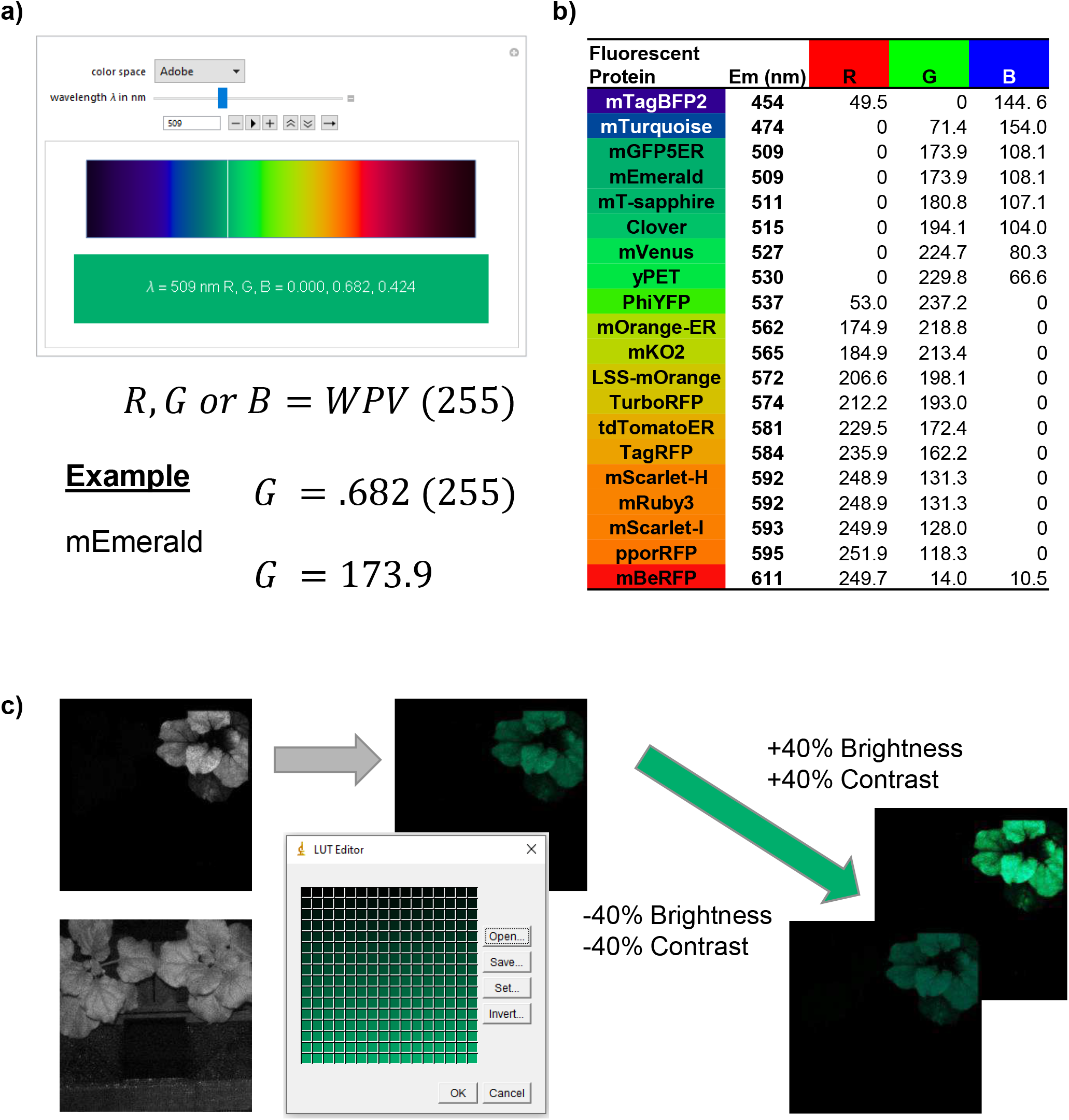
Determination and application of image color. **(a)** Screenshot of the Wolfram Player Colors of the Visible Spectrum plugin for mEmerald peak emission (509) and the formula for the conversion of the Wolfram Player Value (WPV) to red (R), green (G) or blue (B) intensity values. **(b)** Fluorescent protein table that contains peak emission values and the RGB intensity values used to establish the look up tables for pseudo color imaging. **(c)** Example of image processing using the ImageJ software and the application of pseudo color to a FILP image. Includes a screenshot of the ImageJ LUT editor. The example given is the mEmerald image used for the production of Figure 2.

## Acknowledgements

Special thanks to all members of the Center for Agricultural Synthetic Biology at the University of Tennessee for their support as well as lab members Lezlee Dice, Taylor Frazier-Douglas, Cassie Halvorsen, Li Li, Mitra Mazarei, Reginald Millwood, Mary-Anne Nguyen, Alex Pfoetenhaur, Christiano Piasecki, Rebekah Rogers, Yuanhua Shao, Shamira Sultana, and Yongil Yang. We sincerely appreciate the assistance from Richard Sexton and Vilmos Magda at the University of Tennessee Pendergrass library with the 3D printing of the custom plant stand.

This research was developed with funding from the Defense Advanced Research Projects Agency (DARPA) Award No. HR0011-18-2-0049 and Department of Energy (DOE) Grant No. DESC0018347. The views, opinions and/or findings expressed are those of the authors and should not be interpreted as representing the official views or policies of the Department of Defense or the U.S. Government.

## Author Contributions

**CNS, JDB, SCL, TMS**, and **SBR** conceived of the research. **JDB, JAM, MJF, SCL, TMS**, and **CNS** played roles in designing and building the FILP instrument. **SBR, TMS, JHL, SCL**, and **CNS** wrote and prepared the manuscript. **SBR, KAM, HB** and **JHL** were responsible for the organization of figures. **SBR, TMS, JHL, HB, MJS, KAM, MRP, JSL, MJS, AO, SCL** and **EMS** were responsible for the design and construction of all constructs tested, plant care, potato cell culture, protoplast assays, agroinfiltration experiments, confocal microscopy, and FILP imaging and spectroscopy. **MJS** and **SBR** developed the agroinfiltration apparatus and produced the Supplementary video. **HB** and **JWB** were responsible for the conception and writing of the Fluorologger software. **SBR** and **KAM** designed the custom plant stand.

